# Amplicon sequencing of the 16S-ITS-23S rRNA operon with long-read technology for improved phylogenetic classification of uncultured prokaryotes

**DOI:** 10.1101/234690

**Authors:** Joran Martijn, Anders E. Lind, Ian Spiers, Lina Juzokaite, Ignas Bunikis, Olga Vinnere Pettersson, Thijs J.G Ettema

## Abstract

Amplicon sequencing of the 16S rRNA gene is the predominant method to quantify microbial compositions of environmental samples and to discover previously unknown lineages. Its unique structure of interspersed conserved and variable regions is an excellent target for PCR and allows for classification of reads at all taxonomic levels. However, the relatively few phylogenetically informative sites prevent confident phylogenetic placements of novel lineages that are deep branching relative to reference taxa. This problem is exacerbated when only short 16S rRNA gene fragments are sequenced. To resolve their placement, it is common practice to gather more informative sites by combining multiple conserved genes into concatenated datasets. This however requires genomic data which may be obtained through relatively expensive metagenome sequencing and computationally demanding analyses. Here we develop a protocol that amplifies a large part of 16S and 23S rRNA genes within the rRNA operon, including the ITS region, and sequences the amplicons with PacBio long-read technology. We tested our method with a synthetic mock community and developed a read curation pipeline that reduces the overall error rate to 0.18%. Applying our method on four diverse environmental samples, we were able to capture near full-length rRNA operon amplicons from a large diversity of prokaryotes. Phylogenetic trees constructed with these sequences showed an increase in statistical support compared to trees inferred with shorter, Illumina-like sequences using only the 16S rRNA gene (250 bp). Our method is a cost-effective solution to generate high quality, near full-length 16S and 23S rRNA gene sequences from environmental prokaryotes.

## Introduction

The 16S rRNA gene has been used for decades to phylogenetically classify bacteria and archaea (Woese and Fox 1977). The gene excels in this respect because of its universal occurrence, resistance to horizontal gene transfer, and high degree of conservation (Woese 1987, Green and Noller 1997). Highly conserved regions are interspersed with highly variable regions, allowing for phylogenetic classification at all taxonomic levels. In addition, the gene has proven to be an ideal target for studies aiming to quantify the taxonomic composition of microbial communities via high-throughput PCR amplicon sequencing (Doolittle 1999). Primers are usually designed such that they anneal to stretches of conserved sites that flank a variable region, in effect capturing the informative variable region of a large fraction of the microbial community. 16S rRNA amplicon surveys are now a standard method in microbial ecology and has led to important insights into the taxonomic makeup of many different environments. Examples include oceanic waters (Sogin et al. 2006), deep sea sediments (Jorgensen et al. 2012), hot springs (Hou et al. 2013) and the human gut (Turnbaugh et al. 2007).

Despite its many advantages, the 16S rRNA gene is limited in its number of phylogenetically informative sites. 16S rRNA gene-based phylogenies are therefore sensitive to stochastic error and exhibit limited resolution (Brown et al. 2001, Delsuc et al. 2005). Studies often favor large datasets of conserved protein coding genes over the 16S rRNA gene to overcome such error and increase resolution (Brochier et al. 2002, Brown et al. 2001, Wolf et al. 2001, Matte-Tailliez et al. 2002). Another frequently used method is to concatenate 16S rRNA with the larger 23S rRNA gene (Ferla et al. 2013, Zaremba-Niedzwiedzka et al. 2017, Williams et al. 2012). However, both methods require that genome sequences are available for the taxa in question. When investigating communities of previously unexplored environments, these are virtually never available. Though it is now possible to obtain genomic data via metagenomic binning (Albertsen et al. 2013, Alneberg et al. 2014), it requires relatively expensive deep sequencing and computationally demanding metagenome assembly. In addition, genomes acquired via binning often lack rRNA genes (Hugenholtz et al. 2016, Nelson and Mobberley 2017), making it impossible to link genomic bins to environmental lineages observed in 16S rRNA amplicon surveys. One possible solution would be to obtain 16S and 23S rRNA gene sequences simultaneously via PCR amplicon sequencing. This approach would make use of the fact that both genes are neighboring in 67 % of known bacterial and 74 % of all known archaeal genomes (Supplementary Table 1). However, standard high-throughput sequencing methods generate too short sequence reads to efficiently capture such long amplicons.

With the introduction of PacBio’s single-molecule real-time (SMRT) sequencing technology, sequencing long amplicons at moderately high throughput became a possibility. Its initial high error rates (up to 15%) have now substantially been reduced via circular consensus sequencing (ccs). A number of pioneering studies have already used the technology to successfully obtain near full length 16S rRNA gene sequences with low error rates (Schloss et al. 2016, Singer et al. 2016, Wagner et al. 2016). Here we go one step further and obtain near full length 16S and 23S rRNA genes by sequencing amplicons spanning a large part of the rRNA operon. We develop a read curation pipeline that deals with PacBio-specific artefacts, and evaluate error rates with a phylogenetically diverse mock community. We compare our method with classic partial 16S rRNA amplicon sequencing with respect to resolving deep phylogenetic relationships of novel taxa and apply our method to four diverse environmental samples.

## Materials and methods

### Fraction of reference genomes with neighboring 16S and 23S rRNA genes

The fraction of bacterial and archaeal genomes (NCBI RefSeq, 2017-05-19) in which at least one 16S rRNA gene is situated upstream of the 23S rRNA gene (16S-ITS-23S operon size ≤ 6500 bp) was estimated by checking for this gene organization in reference genomes (one representative per species; 12596 bacteria and 364 archaea) with an in-house perl script.

### Primer design

For the forward primer, we used the A519F (5′-CAGCMGCCGCGGTAA-3′) (derived from(Klindworth et al. 2013) primer). It covers a large fraction of the known bacterial and archaeal diversity and has shown robust performance in previous 16S rRNA amplicon studies (Spang et al. 2015, Baker et al. 2016).

For our reverse primer, we designed one in such a manner that it would anneal to the 3′ end of the 23S rRNA gene and covers an as large diversity of bacteria and archaea as possible. First, we took the full, 150,000 nt long SILVA LSURef (release 119) alignment (SILVA_119_LSURef_tax_silva_full_align_trunc.fasta, available at the SILVA archive) and removed all sequences that corresponded to eukaryotes or were shorter than 2200 bp (not counting gaps). We removed all sites with more than 10% gaps (TRIMAL v1.4 -gt 0.90). The archaea are heavily underrepresented in the LSURef 119 database (43822 bacterial entries vs 629 archaeal entries), and any primer based on the current alignment may have a strong bias towards bacteria. To prevent this, and further prevent a bias towards species that have entries for a disproportionate number of strains, we clustered all bacterial entries into 90% OTUs with UCLUST (Edgar 2010) and rebuilt the alignment with one representative sequence per OTU, and supplemented it with all archaeal entries.

We fed this alignment to WEBLOGOv3 (http://weblogo.threeplusone.com/) and ran it in both bit and probability modes with otherwise default settings. The bit mode visualizes highly conserved sites, while the probability mode visualizes the relative occurrence of each base per site. We then used both logos to design candidate primers. They were required (i) anneal to a highly conserved region on the 3′ region of the 23S rRNA gene, (ii) lack degenerate bases in the 3′ end and generally as few degeneracies as possible, (iii) have a predicted melting temperature (T_M_) that is within 5°C of the predicted T_M_ of the forward primer, (iv) have a 3′ terminal G or C to facilitate primer extension and (v) have a low probability of forming homodimers or crossdimers with the forward primer under PCR conditions. Expected TM and probability of primer-dimers were evaluated with Thermo Fisher Scientific’s online MULTI PRIMER ANALYZER.

Taxonomic coverage of all candidate primers was then evaluated with Silva’s TESTPROBE (Quast et al. 2012). In the end, primer ‘U2428R’ (5′-CCRAMCTGTCTCACGACG-3′) was deemed optimal. It covers 98.9% of all bacterial, and 89.5% of all archaeal 23S rRNA gene entries in the SILVA 128 release.

### Mock community

We constructed a ‘mock community’ sample composed of genomic DNA from 38 phylogenetically diverse taxa from bacteria and archaea. To be included, taxa were required to have a complete genome available and have at least one neighboring 16S and 23S rRNA gene (ITS < 1 kbp; for a complete list, see Supplementary Table 2). Genomic DNA for these taxa was ordered from DSMZ and quantified with the Quant-IT PicoGreen dsDNA Assay kit (ThermoFisher) using the FLUOstar Omega microplate reader (BMG Labtech). The genomic DNA samples of all selected taxa were pooled in such a manner that all taxa had an equal amount of gDNA with neighboring 16S and 23S rRNA genes present in the final mock community. For example, the mock would have ten times more *Desulfovibrio gigas* gDNA than to *Bacillus amyloliquefaciens* gDNA, because *D. gigas* encodes a single 16S+23S rRNA gene pair, whereas *B. amyloliquefaciens* encodes 10 pairs. To calculate for each species the volume fraction in the mock, we first converted the measured concentrations in ng/μl to concentrations in ‘operonmol’μl (Supplementary Table 1). An operonmol here is defined as the number of 16S+23S rRNA gene pairs present in a genome multiplied with the molecular weight of the genome. The volume fraction is then calculated by dividing the inverse, μl/operonmol with the sum of μl/operonmol of all taxa.

### Environmental samples

Four environmental samples were used in this study: “P19”, “PM3”, “TNS08”, and “SALA”. P19 is a sediment sample obtained from Radiata Pool, Ngatamariki, New Zealand (Zaremba-Niedzwiedzka et al. 2017). PM3 is a sediment sample taken from 1.25 m below the sea floor using a gravity core at Aarhus Bay, Denmar (Zaremba-Niedzwiedzka et al. 2017, Starnawski et al. 2017). TNS08 is a sediment sample taken from a shallow submarine hydrothermal vent field near Taketomi Island (Zaremba-Niedzwiedzka et al. 2017). SALA is a sample of a black biofilm that was taken at 60m depth in an old silver mine near Sala, Sweden. Detailed descriptions of DNA extractions and further DNA sample cleaning of the samples P19 (or “P1.0019”), PM3 and TNS08 (or “617-1-3”) can be found in Zaremba-Niedzwiedzka et al, 2017. DNA extraction of SALA was done with the FastDNA 50 ml spin kit for soil (MP Biomedicals).

### PCR

We used primers A519F and U2428R to amplify the rRNA operon between position ~520 of the 16S rRNA gene and position ~2430 of the 23S rRNA gene in the mock community and the extracted DNA from the four environmental samples. All PCR reactions were set up with the Q5 High-Fidelity DNA Polymerase kit (New England Biolabs) according to manufacturer’s recommended reaction mix except for a final Q5 concentration of 0.04 U/μl instead of 0.02 U/μl. Each reaction included the Q5 High GC Enhancer and was done with 2 ng of template DNA, unless otherwise specified. Cycling conditions were as follows: denaturation at 98°C for 30 seconds, followed by 30 cycles of amplification (denaturation at 98°C for 10 seconds, annealing at 64°C for 30 seconds, extension at 72°C for 3 minutes and 30 seconds) and a final extension at 72°C for 10 minutes. By default, each sample was amplified in three parallel 50 μl reactions. Deviations from the default: the mock community amplifications were done in 25 μl reactions with 1 ng of template DNA, the TNS08 sample was amplified in two, 34 cycle parallel reactions with 0.2 ng of template, the PM3 sample was amplified in five parallel reactions, and the P19 sample was amplified in nine parallel reactions, of which three were done with 34 cycles. PCR products were cleaned with AMPure XP beads (Beckman Coulter) according to Illumina’s Nextera DNA Library Prep, Clean Up Library protocol (page 15-16). We used a 2:1 PCR product:beads volume ratio, and eluted in ddH_2_O. In addition to removing PCR reagents, bead purification also removes short DNA fragments such as primers, primer-dimers, and potential small, aspecific PCR products. All purified PCR products were then pooled by sample and quantified with the Qubit dsDNA HS (High-Sensitivity) Assay kit (ThermoFisher Scientific).

### PacBio sequencing

Libraries were prepared by ligating SMRTbell adapters onto the PCR products as described in the ‘Procedures & Checklist – 5 kb Template Preparation and Sequencing’ protocol (without the fragmentation step. Each library was loaded onto the SMRT cells with MagBead loading (one library per SMRT cell), and sequenced on the PacBio RSII SMRT DNA Sequencing System with the P6-C4 chemistry and a movie length of 240 minutes. Circular consensus (ccs) reads were generated from the movies with ‘ReadsOfInsert’ protocol, implemented in the SMRT Analysis v2.3.0 (Patch 5).

### Read curation pipeline

For a graphical overview of the curation pipeline that includes explanations on each curation operation, see Figure 1 & Supplementary Figure 1. We started by discarding ccs reads that had an associated predicted read quality (the ‘rq’ tag in the ccs .bam file) of less than 0.99. We further discarded any high quality read that had an internal window of at least 30 bp with an average phred score of lower than 18 with an in-house script. All non-ambiguous base calls amongst the remaining ccs reads with phred score of 0 were changed to ‘N’ with the MOTHUR v.1.37.4 (Schloss et al. 2009) fastq.info function (pacbio=T). The MOTHUR trim.seqs function was then used to discard all reads with 10 or more consecutive identical base calls (maxhomop=10), reads shorter than 3000 bp (minlength=3000) or longer than 5000 bp (maxlength=5000), reads with more than two mismatches to primers (pdiffs=2), and simultaneously trim recognized primer sequences from the starts and ends of reads (keepforward=F). We further used the demultiplexing capacity of trim.seqs to recognize and consecutively remove reads that started and ended with the same primer. Up to this point each ccs read still represents the positive or negative strand, depending on which strand the polymerase initiated the sequencing. In the next step we ‘polarize’ the reads, meaning that after polarization all reads are in the same direction and represent the same strand. To recognize reads derived from opposite strands, we used the --adjustdirection function of MAFFT v7.050b (Katoh and Standley 2013). We detected chimeras *de novo* and subsequently removed them with MOTHUR’s chimera.uchime (reference=self, chunks=80, abskew=1) and remove.seqs, respectively. On all reads that passed we predicted the partial 16S and 23S rRNA genes and their associated internal transcribed spacer (ITS) with RNAMMER v1.2 (Lagesen et al. 2007). RNAMMER was run in bacterial and archaeal modes. Per read, we chose the gene predictions (bacterial or archaeal) with the highest score. Next we preclustered all reads that have both a 16S and a 23S rRNA gene predicted with VSEARCH v2.4.3 at 99% identity level (Rognes et al. 2016) (--cluster_fast, --id 0.99). Next, we generated majority rule consensus reads for each precluster of size 3 or larger. Such preclusters were aligned with MAFFT-QINSI (--kimura 1) and each alignment was consequently used as input for MOTHUR’s consensus.seqs (cutoff=51). Gaps were removed in the resulting consensus, yielding final precluster consensus sequences. Finally, partial 16S and 23S rRNA genes were predicted with RNAMMER as above from both precluster consensus sequences and all reads from clusters of size one and two. All predicted 16S and 23S rRNA genes were used for the phylogenetic tree depicting the environmental diversity, while only predicted 16S and 23S rRNA genes derived from consensus sequences were used for the phylogenetic comparison between Illumina-length and PacBio-length based phylogenies. In total we produced 367 archaeal and 323 bacterial high quality 16S and 23S rRNA consensus sequence pairs. In addition to this, we also generated 11528 non-consensus sequences which were used to examine the total environmental diversity.

**Figure 1.**
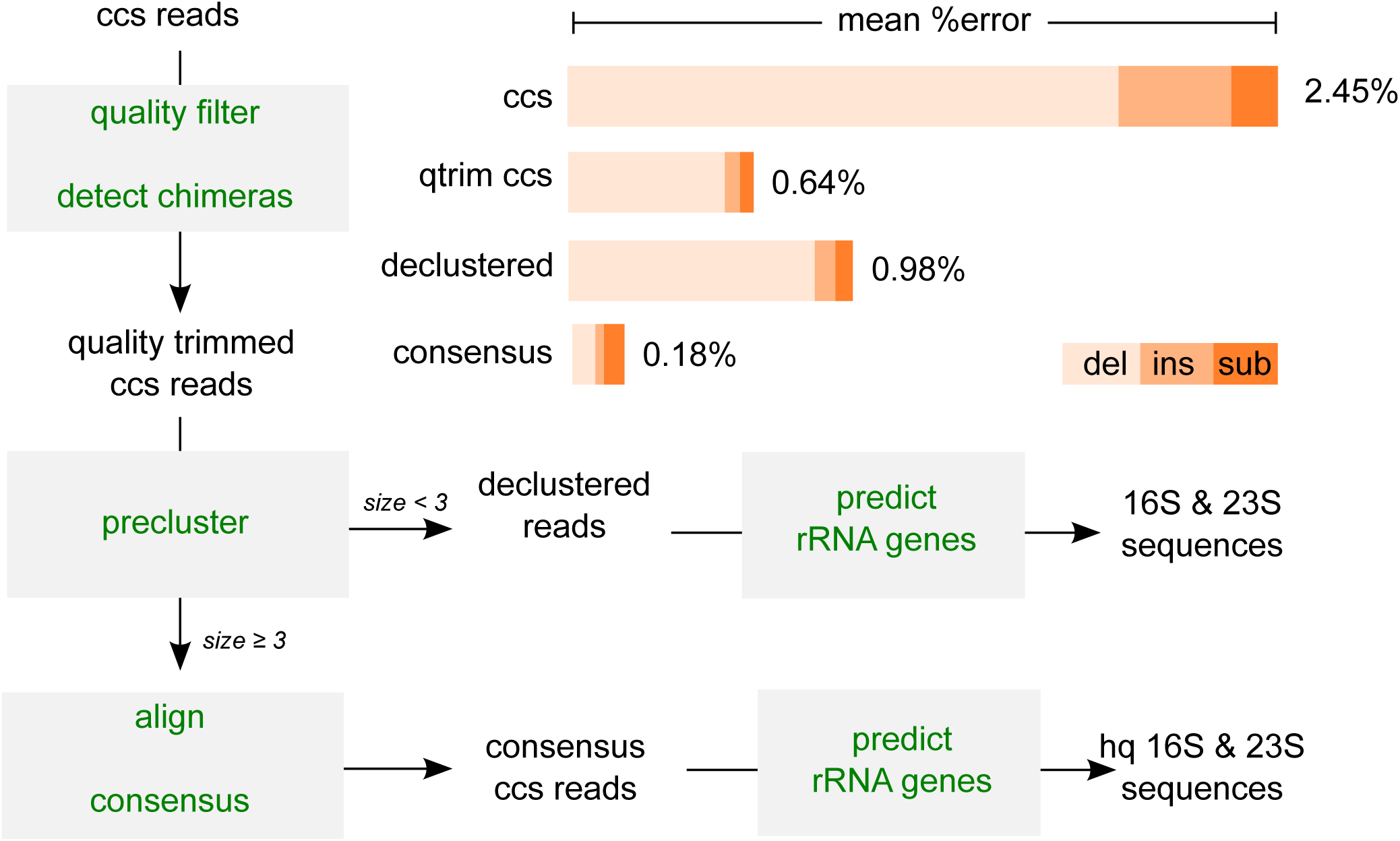
Read curation pipeline and observed mean error rates. For a more detailed overview of the pipeline see Supplementary Figure 1. ‘del’ = deletions, ‘ins’ = insertions, ‘sub’ = substitutions.

### Error rate evaluation

The degree of erroneous base calls was evaluated by comparing the ccs reads derived from the mock community with the reference 16S-ITS-23S sequences at different points in the read curation pipeline (Figure 1). We extracted reference 16S-ITS-23S sequences from the mock community taxa genomes and only included instances smaller than 6000 bp. The reference was further supplemented with 16S-ITS-23S sequences from taxa that were found to contaminate the mock community (*Veillonella parvula* DSM 2008, *Moellerella wisconsensis* ATCC 35017, *Staphylococcus epidermidis* ATCC 12228 and *Streptococcus pneumoniae* R6). For *M. wisconsensis*, the 16S and 23S rRNA genes were encoded on different contigs thus do not necessarily form a 16S-ITS-23S locus. However, since we identified ccs reads with *M. wisconsensis* 16S and 23S rRNA genes, we conclude that they do. We patched the reference 16S and 23S rRNA gene sequences with the ITS sequence from the highest quality *M. wisconsensis* read, and included it in our reference. Ccs reads were compared with the reference by mapping them onto the reference with BLASR v3.1 (-minMatch 15, -maxMatch 20, -bestn 1) (Chaisson and Tesler 2012). Number of substitutions, insertions, deletions were extracted from the CIGAR string and the NM tag in the output SAM file, and used to calculate the overall error rate (sum of substitutions, insertions and deletions divided by the alignment length).

### Phylogenetic analyses

A reference phylogenetics dataset was constructed as follows: all taxa that had 16S (≥ 900 bp) and 23S (≥ 1500 bp) rRNA gene sequences available in SILVA (Pruesse et al. 2007), JGI and NCBI were gathered. Seqeunces were split into to separate Archaea and Bacteria datasets, and the genetic redundancy of the dataset was reduced by grouping the taxa into clusters with VSEARCH (Rognes et al. 2016) (similarity thresholds: Archaea 95%, Bacteria 85%) based on their 16S rRNA gene sequences and keeping only centroid taxa. A lower threshold was used for bacteria to reduce the total number of taxa in the final tree. The 16S and 23S rRNA gene sequences were aligned separately using mafft-qinsi 7.309 (Katoh and Standley 2013) with --maxiterate set to 0, trimmed using trimAL using -gt 0.5 (Capella-Gutiérrez et al. 2009), and a phylogenetic tree was constructed from the concatenated 16S and 23S rRNA genes using IQ-TREE 1.6 beta 4 (Nguyen et al. 2014). Miss-classified taxa (bacteria classified as archaea and *vice versa*) were removed from the dataset. The final reference dataset consisted of 228 archaea, and 618 bacteria.

All high quality pre-cluster sequences were classified as either bacterial or archaeal using MOTHURs classify_seqs function. The sequences were then combined with the appropriate reference dataset (bacterial or archaeal), and the 16S and 23S rRNA sequences were aligned separately using mafft-linsi with --maxiterate set to 0 (Katoh and Standley 2013). Any position in the alignment which was only present in 20 % of all taxa were removed using trimAl 1.4 (Capella-Gutiérrez et al. 2009). Phylogenetic trees were constructed from the 16S and 23S rRNA concatenated alignments using IQ-TREE 1.5.3 (Nguyen et al. 2014) using 1000 ultrafast bootstraps and the GTR+R10 evolutionary model (selected by IQ-TREE’s modeltest).

To compare how our concatenated 16S+23S rRNA gene alignments performed against a more traditional 16S amplicon survey, we took the high-quality preclusters and shortened them to 250 bp using fastx_trimmer (Gordon and Hannon 2010). These 250 bp pseudo-amplicons were again identified as either archaeal or bacterial, and used together with the 16S rRNA gene reference dataset to construct phylogenetic trees in the same way as described above, however the alignments were made using mafft-linsi in iterative mode, and only the 16S rRNA gene dataset was used.

A complete tree utilizing all amplicons, no preclustering performed, and all reference sequences were computed in order to asses any missed diversity due to quality filtering. The sequences were aligned separately for the 16S and 23S sequences using mafft (Katoh and Standley 2013) and trimmed using trimAl with the -gt 0.05 setting. A phylogenetic tree was constructed using IQ-TREE 1.6 Beta (Nguyen et al. 2014) using the -fast option and the GTR+R7 model of evolution. SH-like approximate likelihood ratio test were performed instead of bootstraps.

## Results and discussion

### Read curation pipeline

Here we present a method for generating and sequencing amplicons of approximately 4000 bp containing near full-length 16S and 23S rRNA genes from environmental taxa, including the ITS region. Because the method uses PacBio sequencing technology which exhibits higher error rates compared to Illumina, we developed a read curation pipeline (Figure 1, Supplementary Figure 1) that reduces the mean error rate to an acceptable level. To evaluate the error rate, we applied the method to a synthetic ‘mock community’ composed of the genomic DNA of 38 phylogenetically diverse archaea and bacteria for which complete genomes are available and encode at least one 16S-ITS-23S cluster. In the raw sequence data, we observed a mean error rate of 2.45%. The large majority of errors were deletions (77.7%), followed by insertions (15.8%) and substitutions (6.5%) (Figure 1).

We sought to reduce the error rate by removing reads that are highly erroneous because of several reasons. First, as was observed by Schloss (Schloss et al. 2016), error rates were strongly correlated with the ‘read quality’ values that are calculated by the SMRT analysis software (Supplementary Figure 2). We removed all reads with an associated read quality value of lower than 0.99. Second, a small fraction (1.0%) of high quality reads contained local stretches of consecutive low quality base-calls with an enriched number of errors (Supplementary Figure 3). We therefore removed all reads that contained windows of at least 30 bp with an average Phred score of 18 or lower. Third, we observed a large number of high quality reads (11.3%) that have been previously referred to as ‘siamaeras’ (Hackl et al. 2014). The first half of these reads consist of the expected amplicon, while the second half consists of the reverse complement of the first half, but missing a primer at the breakpoint (Supplementary Figure 4). Siamaeras most likely stem from damaged amplicons that have a long overhang. The overhang forms a hairpin which anneals to the complementary strand. As a result, the SMRTbell adapter is blocked from ligating there during the library preparation, and the read processing software will interpret the concatenation of both strands as a single insert (Supplementary Figure 4). Since siamaeras are about twice the expected read length, they are easily detected and removed by setting a length cutoff (here 5 kbp). However, this does not remove all siamaeras. Some stem from partial rather than full amplicons and may as a result be shorter than the length cutoff. To detect these cases, we use another property of siamaeras: that they start and end with the same primer. We found that the large majority of identified siamaera’s start and end with the reverse primer, implying that choice of primers affects siamaera formation. More analyses are necessary to resolve this issue. Fourth, since our method is PCR based, we need to detect and remove chimeras. Chimeras are formed when incomplete extensions from one locus anneal to another locus, either within the same genome or on another genome. A frequently used method to detect chimeras uses a *de novo* approach (Edgar et al. 2011). A query read is split into four equally sized ‘chunks’ and compared against more abundant reads of the same dataset to find two candidate parents. If a model chimera constructed from the two parents is more similar to the query than each original parent, and at least one of the parent reads is at least two times more abundant than the query, the read is deemed chimeric. The method thus assumes that chimeric amplicons are less abundant than non-chimeric amplicons. It works well for Illumina data, but needs to be adjusted for PacBio data. The lower throughput, longer reads, and higher error rate mean that virtually all reads are unique and thus have an abundance of one. As a result, a chimera will have the same abundance as a parent and will not be detected. In addition, because only four chunks are used by default, chimeras with breakpoints in the first or last ~1000 bp may be missed. Thus, to account for the nature of the PacBio data we used an abundance ratio threshold of one, and increased the number of chunks to 80. Finally, since some reads may stem from loci other than 16S-ITS-23S, we removed all reads that did not contain both genes. After these operations, the error rate was reduced to 0.64% (Figure 1). Deletions were still the most prominent type of error (84.4%), followed by insertions (8.3%) and substitutions (7.2%) (Figure 1).

Though a substantial improvement, we sought to reduce the error rate even further. We argued that if the remaining read errors are still randomly distributed over the reads, then reads originating from the same locus or the same genome will have errors at different positions. As a result, true base-calls would outnumber erroneous base-calls per site, and their consensus would have a reduced error rate. We grouped reads putatively originating from the same genome by clustering the reads at a 99% identity threshold. This is similar to the preclustering method of Schloss (Schloss et al. 2009) that constructs a final set of high quality reads by selecting the most abundant read per 99% precluster. Since there is virtually no abundance information of longer PacBio reads, we preferred a consensus based method. When only considering preclusters with at least three reads, the error rate dropped to 0.18%. Deletions were no longer as prominent (44.9%), now more comparable to insertions (17.5%) and substitutions (37.5%) (Figure 1). We deem these consensus reads high quality and could be used as references for future studies. On the other hand, when considering reads derived from preclusters of size one or two, the mean error rate was 0.98% (Figure 1). Deletions represented the large majority of the errors (86.8%), followed by insertions (6.9%) and substitutions (6.3%). Since deletions translate into gaps, and phylogenetic analyses are generally robust to gaps (missing data; (Philippe et al. 2004), we deem these reads still useful for the phylogenetic identification and placement of environmental taxa.

It should be noted that our consensus method works well for our mock community because all taxa are phylogenetically distinct. For environmental samples that may contain a certain degree of strain microdiversity, 99% preclusters will most likely contain reads originating from various strains in addition to from various loci of the same strain. The consensus sequences will thus represent not the 16S-ITS-23S sequence of one strain, but of multiple closely related strains.

### Phylogenetic analyses

Combining the 16S and the 23S rRNA gene in phylogenetic analysis is a clear advantage to using only the 16S rRNA gene as the number of informative sites increases and thus the phylogenetic signal becomes stronger (Ferla et al. 2013, Zaremba-Niedzwiedzka et al. 2017, Williams et al. 2012). To compare our method with standard 16S rRNA amplicon sequencing with respect to phylogenetic classification, we constructed the following phylogenetic datasets: ‘16S+23S’, a concatenation of full-length reference 16S and 23S rRNA genes, supplemented with the processed full-length PacBio precluster consensus reads, and ‘16S’, containing full length reference 16S rRNA genes, supplemented with shortened processed PacBio precluster consensus reads (250 bp) to simulate Illumina type lengths. For each dataset, a bacterial and archaeal version was constructed. We then inferred maximum likelihood phylogenies for all datasets, and compared the overall topology and statistical support of the obtained trees between 16S and 16S+23S datasets. We observed topological changes and an increase in statistical support for many of the branches of the tree, regardless of using bacterial or archaeal datasets (Figure 2, Supplementary Figure 5). An example of a major topological shift is the sequence ‘PM3_PC_778’, which branches as a sister species to the Heimdallarchaeota in the 16S+23S tree, while it branches as a sister to the Bathyarchaeota in the 16S tree (Supplementery Figure 6). Another example is a group of sequences that constitute a long branch within Actinobacteria with low bootstrap support in the 16S tree, while branching within the recently described Candidate Phyla Radiation (CPR; (Hug et al. 2016)) with maximum support in the 16S+23S tree (Supplementary Figure 7). This suggests that these sequences could not have been identified as members of CPR by standard 16S amplicon assays.

**Figure 2.**
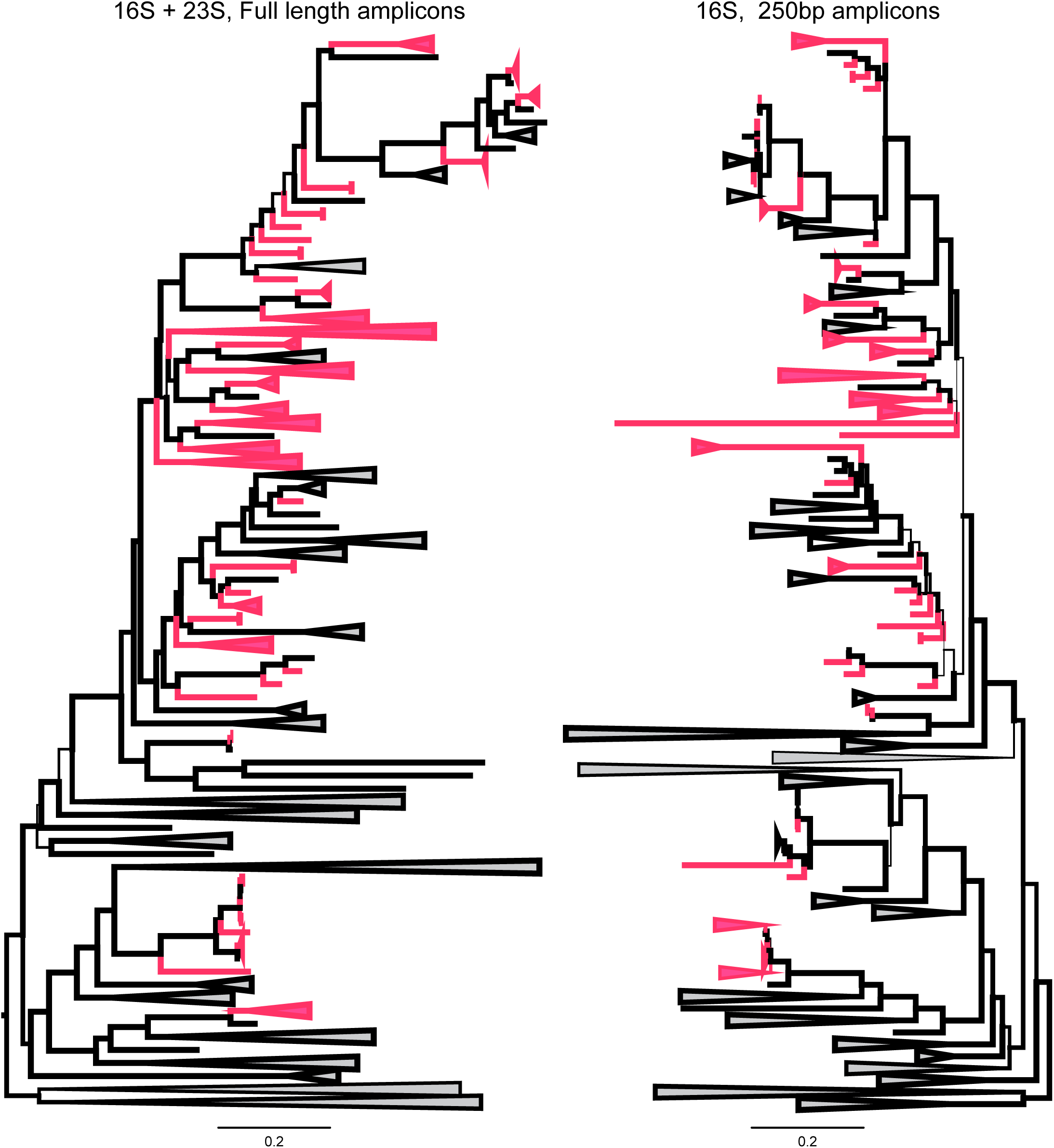
Phylogenetic trees of the archaea using either a concatenated 16S and 23S sequence or 16S only. Left side shows the 16S-23S dataset and the right side the 16S dataset using 250 bp amplicons. Branch bootstrap support is indicated by the thickness of the branches. Branches containing only amplicon sequences are shown in pink and reference sequences in grey. The root of the tree was set using DPANN as an outgroup. Scale bar represents substitutions per site.

The read curation pipeline requires preclusters to have at least three members to generate a high quality consensus read. As a result, rare yet phylogenetically novel lineages may be missed when only considering consensus reads. Reads of smaller preclusters may have higher error rates, but because the errors stem largely from deletions, they are still useful for identification of such lineages. Indeed, when we add these reads to the phylogenetic analysis, we observed several lineages that were missed by the consensus reads (Supplementary Figure 8). Examples include previously undetected members of DPANN archaea, Alphaproteobacteria, Firmicutes and Cyanobacteria. It should be noted however that most of the diversity captured by the reads of smaller preclusters overlapped with that captured by the consensus reads. In addition, since these reads exhibit somewhat higher error rates, one should be aware that any microdiversity observed in the tree associated with these reads may stem from errors rather than biological diversity.

### Expanding available reference databases

The 16S rRNA gene has long been a key tool in phylogenetic and ecological analyses (Lane et al. 1985, Weisburg et al. 1991). Previous studies have used PCR primers targeting the 16S gene to asses microbial diversity from various environments (Turnbaugh et al. 2007, Hou et al. 2013), and thus the amount of available (full or partial) 16S rRNA gene sequences in public databases is large. Though amplicon studies targeting the 23S rRNA gene have been done in the past (Zimmermann et al. 2005, Hunt et al. 2006), the retrieval of 23S rRNA gene sequences relies mostly on genome sequencing projects. This is especially visible for the archaeal domain, for which only 1352 LSU rRNA sequences are available in the SILVA database (release 128; (Pruesse et al. 2007)). Here, we generate a total of 367 high quality archaeal 23S rRNA gene sequences, which would represent a 27% increase. Our method could thus lead to an overall better representation of archaeal diversity in 23S rRNA gene databases.

### Internal transcribed spacer (ITS)

The method presented here provides several advantages over existing environmental rRNA gene survey methods, due to the fact that we are able to get sequence information from a large part of the 16S-ITS-23S cluster. Interestingly, apart from the rRNA genes themselves, we are also able to sequence the internal transcribed spacer (ITS). The ITS, which tend to be faster evolving compared to the rRNA genes (Barry et al. 1991), can be used in conjuction with the 16S and 23S rRNA gene to further increase the resolution of the classification when characterizing closely related species. The use of the ITS region as a taxonomical marker has been previously used in fungi, where the 18S rRNA gene alone does not provide sufficient taxonomic information (Nilsson et al. 2009).

## Acknowledgements

This work is supported by grants of the European Research Council (ERC Starting grant 310039-PUZZLE_CELL), the Swedish Foundation for Strategic Research (SSF-FFL5) and the Swedish Research Council (VR grant 2015-04959) awarded to T.J.G.E. We thank F. Homa, K. Katoh, M. Hammond, J. Vosseberg, C. Bergin, A.M. Divne and the technical support of New England Biolabs and Pacific Biosciences for useful advice and insightful discussions.

## Author contributions

T.J.G.E. conceived the study. J.M. and I.S. designed the primers. J.M., I.S. and L.J. designed and constructed the mock community, and developed the PCR protocol. L.J. extracted DNA from the environmental samples. I.B. and I.V.P. performed PacBio sequencing. J.M. developed the read curation pipeline. A.E.L. performed the phylogenetic analyses. J.M., A.E.L. and T.J.G.E. analyzed and interpreted the results and wrote the manuscript. All authors edited and approved the manuscript.

## References

Albertsen, Mads, Philip Hugenholtz, Adam Skarshewski, Kåre L. Nielsen, Gene W. Tyson, and Per H. Nielsen. 2013. “Genome sequences of rare, uncultured bacteria obtained by differential coverage binning of multiple metagenomes.” Nature Biotechnology 31:533–538.

Alneberg, Johannes, Brynjar Smári Bjarnason, Ino de Bruijn, Melanie Schirmer, Joshua Quick, Umer Z. Ijaz, Leo Lahti, Nicholas J. Loman, Anders F. Andersson, and Christopher Quince. 2014. “Binning metagenomic contigs by coverage and composition.” Nature Methods 11:1144–1146.

Baker, Brett J., Jimmy H. Saw, Anders E. Lind, Cassandre Sara Lazar, Kai-Uwe Hinrichs, Andreas P. Teske, and Thijs J. G. Ettema. 2016. “Genomic inference of the metabolism of cosmopolitan subsurface Archaea, Hadesarchaea.” Nature Microbiology 1:16002.

Barry, Tom, Gerard Colleran, Maura Glennon, LK Dunican, and F Gannon. 1991. “The 16s/23s ribosomal spacer region as a target for DNA probes to identify eubacteria.” Genome Research 1 (1):51–56.

Brochier, Céline, Eric Bapteste, David Moreira, and Hervé Philippe. 2002. “Eubacterial phylogeny based on translational apparatus proteins.” Trends in Genetics 18:1–5.

Brown, James R., Christophe J. Douady, Michael J. Italia, William E. Marshall, and Michael J. Stanhope. 2001. “Universal trees based on large combined protein sequence data sets.” Nature Genetics 28:281–285.

Capella-Gutiérrez, Salvador, José M. Silla-Martínez, and Toni Gabaldón. 2009. “trimAl: a tool for automated alignment trimming in large-scale phylogenetic analyses.” Bioinformatics 25:1972–1973.

Chaisson, Mark J., and Glenn Tesler. 2012. “Mapping single molecule sequencing reads using basic local alignment with successive refinement (BLASR): application and theory.” BMC Bioinformatics 13:238.

Delsuc, Frédéric, Henner Brinkmann, and Hervé Philippe. 2005. “Phylogenomics and the reconstruction of the tree of life.” Nature Reviews Genetics 6:361–375.

Doolittle W Ford. 1999. “Phylogenetic classification and the universal tree.” Science 284 (5423):2124–2128.

Edgar, Robert C, Brian J Haas, Jose C Clemente, Christopher Quince, and Rob Knight. 2011. “UCHIME improves sensitivity and speed of chimera detection.” Bioinformatics 27 (16):2194–2200.

Edgar, Robert C. 2010. “Search and clustering orders of magnitude faster than BLAST.” Bioinformatics 26:2460–2461.

Ferla, Matteo P., J. Cameron Thrash, Stephen J. Giovannoni, and Wayne M. Patrick. 2013. “New rRNA Gene-Based Phylogenies of the Alphaproteobacteria Provide Perspective on Major Groups, Mitochondrial Ancestry and Phylogenetic Instability.” PLoS ONE 8:e83383.

Gordon, A, and GJ Hannon. 2010. “Fastx-toolkit.” FASTQ/A short-reads preprocessing tools (unpublished) http://hannonlab.cshl.edu/fastx_toolkit.

Green, Rachel, and Harry F Noller. 1997. “Ribosomes and translation.” Annual review of biochemistry 66 (1):679–716.

Hackl, Thomas, Rainer Hedrich, Jörg Schultz, and Frank Förster. 2014. “proovread : large-scale high-accuracy PacBio correction through iterative short read consensus.” Bioinformatics 30:3004–3011.

Hou, Weiguo, Shang Wang, Hailiang Dong,Hongchen Jiang, Brandon R Briggs, Joseph P Peacock, Qiuyuan Huang, Liuqin Huang,Geng Wu, and Xiaoyang Zhi. 2013. “A comprehensive census of microbial diversity in hot springs of Tengchong, Yunnan Province China using 16S rRNA gene pyrosequencing.” PloS one 8 (1):e53350.

Hug, Laura A., Brett J. Baker, Karthik Anantharaman, Christopher T. Brown, Alexander J. Probst, Cindy J. Castelle, Cristina N. Butterfield, Alex W. Hernsdorf, Yuki Amano, Kotaro Ise,Yohey Suzuki, Natasha Dudek,David A. Relman, Kari M. Finstad, Ronald Amundson, Brian C. Thomas, and Jillian F. Banfield. 2016. “A new view of the tree of life.” Nature Microbiology 1:16048.

Hugenholtz, Philip, Adam Skarshewski, and Donovan H. Parks. 2016. “Genome-Based Microbial Taxonomy Coming of Age.” Cold Spring Harbor Perspectives in Biology 8:a018085.

Hunt, Dana E, Vanja Klepac-Ceraj, Silvia G Acinas, Clement Gautier, Stefan Bertilsson, and Martin F Polz. 2006. “Evaluation of 23S rRNA PCR primers for use in phylogenetic studies of bacterial diversity.” Applied and environmental microbiology 72 (3):2221–2225.

Jorgensen, Steffen Leth, Bjarte Hannisdal, Anders Lanzén, Tamara Baumberger, Kristin Flesland,Rita Fonseca, Lise Øvreås, Ida H. Steen, Ingunn H. Thorseth, Rolf B. Pedersen, and Christa Schleper. 2012. “Correlating microbial community profiles with geochemical data in highly stratified sediments from the Arctic Mid-Ocean Ridge.” Proceedings of the National Academy of Sciences 109:E2846–E2855.

Katoh, Kazutaka, and Daron M. Standley. 2013. “MAFFT Multiple Sequence Alignment Software Version 7: Improvements in Performance and Usability.” Molecular Biology and Evolution 30:772–780.

Klindworth, Anna, Elmar Pruesse, Timmy Schweer, Jörg Peplies, Christian Quast, Matthias Horn, and Frank Oliver Glöckner. 2013. “Evaluation of general 16S ribosomal RNA gene PCR primers for classical and next-generation sequencing-based diversity studies.” Nucleic Acids Research 41:e1–e1.

Lagesen, Karin, Peter Hallin, Einar Andreas Rødland, Hans-Henrik Stærfeldt, Torbjørn Rognes, and David W. Ussery. 2007. “RNAmmer: consistent and rapid annotation of ribosomal RNA genes.” Nucleic Acids Research 35:3100–3108.

Lane, D. J., B. Pace, G. J. Olsen, D. A. Stahl, M. L. Sogin, and N. R. Pace. 1985. “Rapid determination of 16S ribosomal RNA sequences for phylogenetic analyses.” Proceedings of the National Academy of Sciences 82:6955–6959.

Matte-Tailliez, Oriane, Céline Brochier, Patrick Forterre, and Hervé Philippe. 2002. “Archaeal phylogeny based on ribosomal proteins.” Molecular biology and evolution 19 (5):631–639.

Nelson, William C., and Jennifer M. Mobberley. 2017. Biases in genome reconstruction from metagenomic data.

Peer J Preprints. Nguyen, Lam-Tung, Heiko A Schmidt, Arndt von Haeseler, and Bui Quang Minh. 2014. “IQ-TREE: a fast and effective stochastic algorithm for estimating maximum-likelihood phylogenies.” Molecular biology and evolution 32 (1):268–274.

Nilsson Rolf Henrik, Martin Ryberg,Kessy Abarenkov, Elisabet Sjökvist, and Erik Kristiansson. 2009. “The ITS region as a target for characterization of fungal communities using emerging sequencing technologies.” FEMS Microbiology Letters 296 (1):97–101.

Philippe, Hervé, Elizabeth A. Snell, Eric Bapteste, Philippe Lopez, Peter W. H. Holland, and Didier Casane. 2004. “Phylogenomics of Eukaryotes: Impact of Missing Data on Large Alignments.” Molecular Biology and Evolution 21:1740–1752.

Pruesse, Elmar, Christian Quast, Katrin Knittel, Bernhard M Fuchs, Wolfgang Ludwig, Jörg Peplies, and Frank Oliver Glöckner. 2007. “SILVA: a comprehensive online resource for quality checked and aligned ribosomal RNA sequence data compatible with ARB.” Nucleic acids research 35 (21):7188–7196.

Quast, C., E. Pruesse, P. Yilmaz, J. Gerken, T. Schweer, P. Yarza, J. Peplies, and F. O. Glockner. 2012. “The SILVA ribosomal RNA gene database project: improved data processing and web-based tools.” Nucleic Acids Research 41:D590–D596.

Rognes, Torbjørn, Tomáš Flouri, Ben Nichols, Christopher Quince, and Frédéric Mahé. 2016. “VSEARCH: a versatile open source tool for metagenomics.” PeerJ 4:e2584.

Schloss, Patrick D., Matthew L. Jenior, Charles C. Koumpouras, Sarah L. Westcott, and Sarah K. Highlander. 2016. “Sequencing 16S rRNA gene fragments using the PacBio SMRT DNA sequencing system.” PeerJ 4:e1869.

Schloss, Patrick D., Sarah L. Westcott, Thomas Ryabin, Justine R. Hall, Martin Hartmann, Emily B. Hollister, Ryan A. Lesniewski, Brian B. Oakley, Donovan H. Parks, Courtney J. Robinson, Jason W. Sahl, Blaz Stres, Gerhard G. Thallinger, David J. Van Horn, and Carolyn F. Weber. 2009. “Introducing mothur: Open-Source, Platform-Independent, Community-Supported Software for Describing and Comparing Microbial Communities.” Applied and Environmental Microbiology 75:7537–7541.

Singer, Esther, Brian Bushnell, Devin Coleman-Derr, Brett Bowman, Robert M. Bowers, Asaf Levy, Esther A. Gies, Jan-Fang Cheng, Alex Copeland, Hans-Peter Klenk, Steven J. Hallam, Philip Hugenholtz, Susannah G. Tringe, and Tanja Woyke. 2016. “High-resolution phylogenetic microbial community profiling.” The ISME Journal 10:2020–2032.

Sogin, Mitchell L., Hilary G. Morrison, Julie A. Huber, David Mark Welch, Susan M. Huse, Phillip R. Neal, Jesus M. Arrieta, and Gerhard J. Herndl. 2006. “Microbial diversity in the deep sea and the underexplored “rare biosphere”.” Proceedings of the National Academy of Sciences 103:12115–12120.

Spang, Anja, Jimmy H. Saw, Steffen L. Jørgensen, Katarzyna Zaremba-Niedzwiedzka, Joran Martijn, Anders E. Lind, Roel van Eijk, Christa Schleper, Lionel Guy, and Thijs J. G. Ettema. 2015. “Complex archaea that bridge the gap between prokaryotes and eukaryotes.” Nature 521:173–179.

Starnawski, Piotr, Thomas Bataillon, Thijs JG Ettema, Lara M Jochum, Lars Schreiber, Xihan Chen, Mark A Lever, Martin F Polz, Bo B Jørgensen, and Andreas Schramm. 2017. “Microbial community assembly and evolution in subseafloor sediment.” Proceedings of the National Academy of Sciences 114 (11):2940–2945.

Turnbaugh, Peter J., Ruth E. Ley, Micah Hamady, Claire Fraser-Liggett, Rob Knight, and Jeffrey I. Gordon. 2007. “The human microbiome project: exploring the microbial part of ourselves in a changing world.” Nature 449:804–810.

Wagner, Josef, Paul Coupland, Hilary P. Browne, Trevor D. Lawley, Suzanna C. Francis, and Julian Parkhill. 2016. “Evaluation of PacBio sequencing for full-length bacterial 16S rRNA gene classification.” BMC Microbiology 16:274.

Weisburg, William G, Susan M Barns, Dale A Pelletier, and David J Lane. 1991. “16S ribosomal DNA amplification for phylogenetic study.” Journal of bacteriology 173 (2):697–703.

Williams, Tom A., Peter G. Foster, Tom M. W. Nye, Cymon J. Cox, and T. Martin Embley. 2012. “A congruent phylogenomic signal places eukaryotes within the Archaea.” Proc. R. Soc. B 279:4870–4879.

Woese, Carl R. 1987. “Bacterial evolution.” Microbiological reviews 51 (2):221.

Woese, Carl R., and George E. Fox. 1977. “Phylogenetic structure of the prokaryotic domain: The primary kingdoms.” Proceedings of the National Academy of Sciences 74:5088–5090.

Wolf, Yuri I., Igor B. Rogozin, Nick V. Grishin, Roman L. Tatusov, and Eugene V. Koonin. 2001. “Genome trees constructed using five different approaches suggest new major bacterial clades.” BMC Evolutionary Biology 1:8.

Zaremba-Niedzwiedzka, Katarzyna, Eva F. Caceres, Jimmy H. Saw, Disa Bäckström, Lina Juzokaite, Emmelien Vancaester,Kiley W. Seitz, Karthik Anantharaman, Piotr Starnawski,Kasper U. Kjeldsen, Matthew B. Stott, Takuro Nunoura, Jillian F. Banfield, Andreas Schramm, Brett J. Baker, Anja Spang, and Thijs J. G. Ettema. 2017. “Asgard archaea illuminate the origin of eukaryotic cellular complexity.” Nature 541:353–358.

Zimmermann, J, JM Gonzalez, C Saiz-Jimenez, and W Ludwig. 2005. “Detection and phylogenetic relationships of highly diverse uncultured acidobacterial communities in Altamira Cave using 23S rRNA sequence analyses.” Geomicrobiology Journal 22 (7-8):379–388.

